# Cotranslational folding of alkaline phosphatase in the periplasm of *Escherichia coli*

**DOI:** 10.1101/2020.07.06.189464

**Authors:** Rageia Elfageih, Alexandros Karyolaimos, Grant Kemp, Jan-Willem de Gier, Gunnar von Heijne, Renuka Kudva

## Abstract

Cotranslational protein folding studies using Force Profile Analysis, a method where the SecM translational arrest peptide is used to detect folding-induced forces acting on the nascent polypeptide, have so far been limited mainly to small domains of cytosolic proteins that fold in close proximity to the translating ribosome. In this study, we investigate the cotranslational folding of the periplasmic, disulfide bond-containing *E. coli* protein alkaline phosphatase (PhoA) in a wild-type strain background and a strain background devoid of the periplasmic thiol:disulfide interchange protein DsbA. We find that folding-induced forces can be transmitted *via* the nascent chain from the periplasm to the polypeptide transferase center in the ribosome, a distance of ~160 Å, and that PhoA appears to fold cotranslationally *via* at least two disulfide-stabilized folding intermediates. Thus, Force Profile Analysis can be used to study cotranslational folding of proteins in an extra-cytosolic compartment, like the periplasm.

## Introduction

Protein secretion across the inner membrane in the Gram-negative bacterium *Escherichia coli* has been classically thought to proceed post-translationally, orchestrated by an interplay between the signal peptide of the protein, cytoplasmic chaperones such as Trigger Factor and SecB, the ATPase SecA, and the SecYEG translocon ^1–17^.

Post-translational export commences when SecA recognizes a secretory protein *via* its signal sequence, and targets it to the inner membrane ^18–22^. Folding *en route* to the membrane is prevented by sequence-specific motifs within the protein itself ^23^, and by interactions with cytoplasmic chaperones such as SecB ^12,24,25^. The unfolded state enables the protein to be exported *via* the SecYEG translocon ^26^, driven by the SecA ATPase activity and by the proton motive force ^27–31^. The signal sequence of the protein is then cleaved off by signal peptidase I (LepB), after most of the protein is completely translocated across the inner membrane into the periplasm ^32,33^. Once in the oxidizing environment of the periplasm, the protein commences folding, often involving the formation of disulfide bridges catalyzed by the periplasmic Disulfide bond (Dsb) system ^34–37^.

The existing view that secretion of proteins across the inner membrane occurs post-translationally has been recently challenged by experiments demonstrating that SecA can associate with ribosomes and ribosome-bound nascent chains during translation ^38–42^. These studies suggest that proteins may also be targeted and secreted cotranslationally, and are consistent with earlier findings that signal peptides of certain periplasmic proteins are cleaved before the proteins are fully exported ^32^. Moreover, it has been found that translation intermediates of alkaline phosphatase (PhoA) can form complexes with the periplasmic thiol:disulfide interchange protein DsbA, indicating that a population of PhoA can fold cotranslationally, as assayed by disulfide-bond formation ^43^.

Force-profile analysis (FPA) is a recently developed method that has been used to study cotranslational protein folding of cytoplasmic proteins, both *in vitro* and *in vivo* ^44–53^. FPA takes advantage of the sensitivity of the SecM-family of translational arrest peptides (APs) to pulling forces acting on the nascent chain: the higher the pulling force, the less efficient is the translational stall induced by the AP ^54,55^. Many cotranslational processes, including protein folding, can generate force on the nascent chain, and are hence amenable to FPA. Here, we ask whether FPA can be used to study cotranslational folding in the periplasm of *E. coli*, *i.e.*, when proteins fold ≳160 Å away from the peptidyl transferase center (PTC) in the 50S subunit of the ribosome, where the force exerted on the nascent chain is sensed by the AP. Specifically, we have analyzed the cotranslational folding of PhoA. Our results suggest the existence of at least two disulfide-stabilized, cotranslational folding intermediates that form when ~330 and ~360 residues of mature PhoA have emerged into the periplasm.

## Results and Discussion

### FPA as a tool to study protein folding in the periplasm

FPA is based on the principle that cotranslational folding of a protein domain fused an AP can generate a pulling force on the AP that reduces its stalling efficiency ^44–53^. To study the cotranslational folding of PhoA by FPA, different lengths of PhoA were fused to the seventeen amino-acids long *E. coli* SecM AP *via* a four amino acid long glycine-serine linker and a hemagglutinin tag (HA), Fig. 1A, B; the length *N* of each construct is calculated from the N-terminal end of PhoA (including the signal peptide) to the last amino acid in the AP. The thirty amino-acids long linker sequence (serine-glycine linker + HA tag + AP) was kept constant for all constructs, to reduce background noise in the force profile stemming from interactions of the elongating chain with the ribosome exit tunnel. A 77-residue long C-terminal tail derived from the *E. coli* protein LacZ was engineered at the C terminus of the AP to ensure good separation of the arrested (*A*) and full-length (*FL*) forms of the protein by SDS-PAGE, Fig. 1C. Since PhoA has a cleavable signal peptide, the possible appearance of both cleaved and uncleaved forms could complicate interpretation of the FPA data due to multiple protein species on SDS-PAGE. To avoid this, we used a variant of PhoA with a point mutation (A_21_P) in the signal peptide that renders it uncleavable, yet does not interfere with export of PhoA into the periplasm ^32,56^.

**Figure 1.**
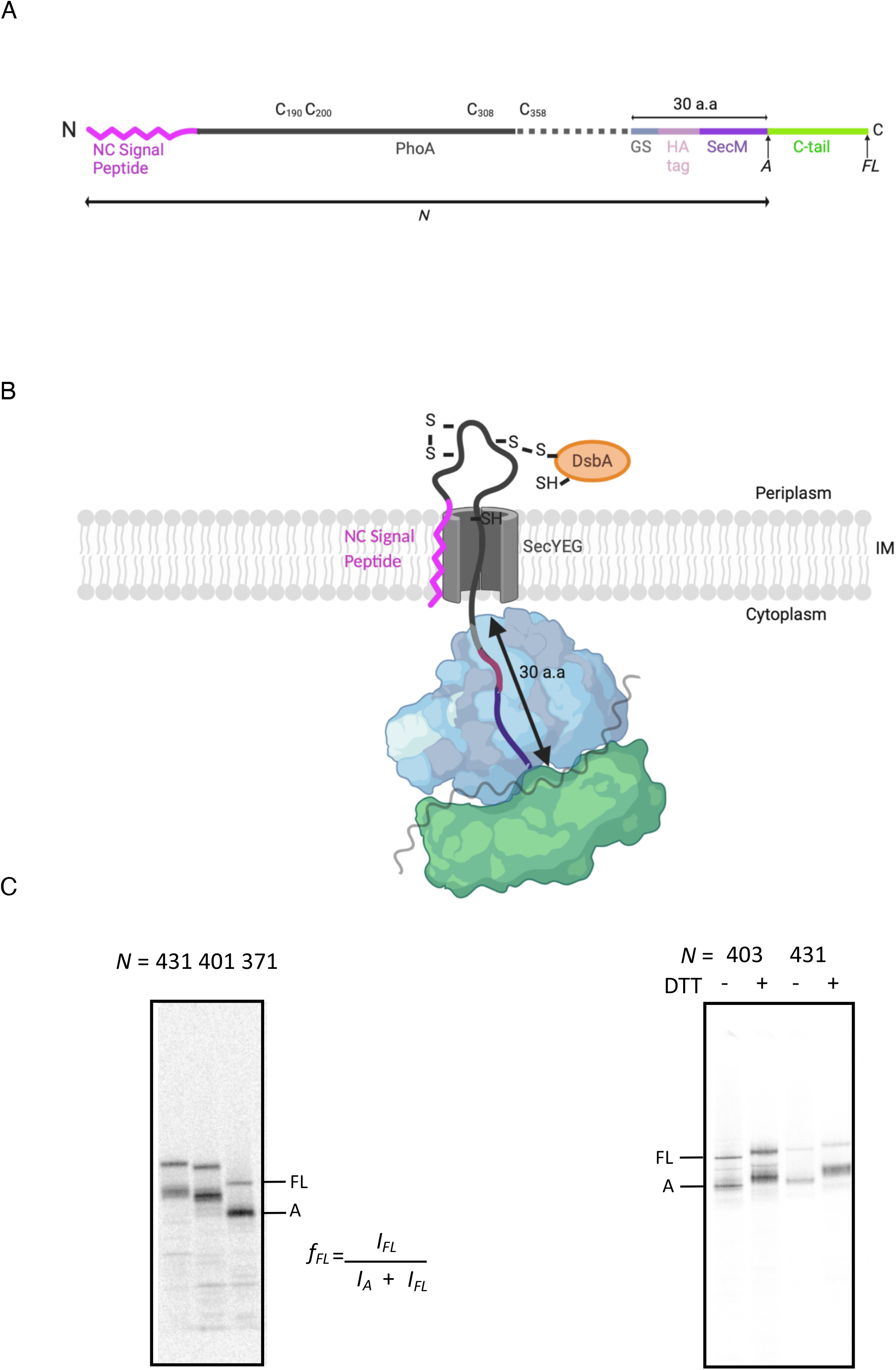
(A) Diagrammatic representation of the PhoA constructs. Different lengths of PhoA with a non-cleavable signal peptide were fused to the *E. coli* SecM arrest peptide (*Ec*SecM) *via* a GS linker and an HA tag (a comprehensive list of sequences for all constructs is included as Supplementary Information). The thirty amino-acid long linker sequence (serine-glycine linker + HA tag + AP) was kept constant for all constructs. A 77 amino acid long C-terminal tail was added at the C-terminal end of the AP to allow for resolution of arrested *A* and full-length *FL* protein products by SDS-PAGE. The different lengths of PhoA were generated by truncating from the C-terminal end of the PhoA coding region. (B) A schematic of cotranslational translocation of PhoA with the non-cleavable signal sequence docked in the membrane and the mature domain translocated into the periplasm where it can form disulfide bonds upon interacting with DsbA. (C) Autoradiographs of three different constructs of SecM-stalled PhoA obtained after radioactive pulse-labelling *in vivo* and SDS-PAGE. The relative amounts of *A* and *FL* were estimated by quantification of the protein bands in the autoradiographs, and the fraction full-length was calculated as *f*_*FL*_=*I*_*FL*_/(*I*_*A*_+*I*_*FL*_). *f*_*FL*_ serves as a proxy for the force generated by cotranslational folding of the mature domain of PhoA in the periplasm. (D) Autoradiographs of constructs *N* = 403 and *N* = 431 demonstrate a difference in protein migration on SDS-PAGE ±DTT. DTT was either excluded or included in the Laemmli buffer that was used to elute the immune-precipitated proteins after pulse-labelling. The differences in migration reflect the formation of disulfide bonds in the absence of DTT, which are reduced in its presence. The schematics in panels A and B were created using BioRender.com.

All constructs were expressed in *E. coli* MC1061 or in MC1061Δ*dsbA*. Proteins were labeled for 2 min. with [^35^S] methionine, followed by immune-precipitation against the HA-tag and separation by SDS-PAGE. As a proxy for the pulling force acting on the nascent chain, the fraction full-length protein (*f*_*FL*_) was calculated for each *N* as the ratio between the intensity of the *FL* protein band and the sum of the intensities of the *A* and *FL* protein bands (Figure 1C). *f*_*FL*_ values were plotted against their corresponding *N* values to obtain a force profile (FP) for PhoA. Both the *A* and *FL* PhoA protein products for lengths *N* = 403 and *N* = 431 showed a difference in their migration on SDS-PAGE in the presence or absence of the reductant 1,4-dithiothreitol (DTT), Fig. 1D, indicating disulfide bond formation and therefore translocation into the periplasm, as depicted in Fig. 1B.

### The PhoA force profile reveals two disulfide bond-stabilized folding intermediates

The PhoA FP obtained in wild-type *E. coli*, Fig. 2A, reveals multiple peaks with maxima at *N* = 311, 403-407, and 431 residues. In addition, the *f*_*FL*_ value for the *N* = 555 construct, in which the entire PhoA domain has been translocated into the periplasm and is connected *via* an 84-residue long linker to the PTC, is also relatively high.

**Figure 2.**
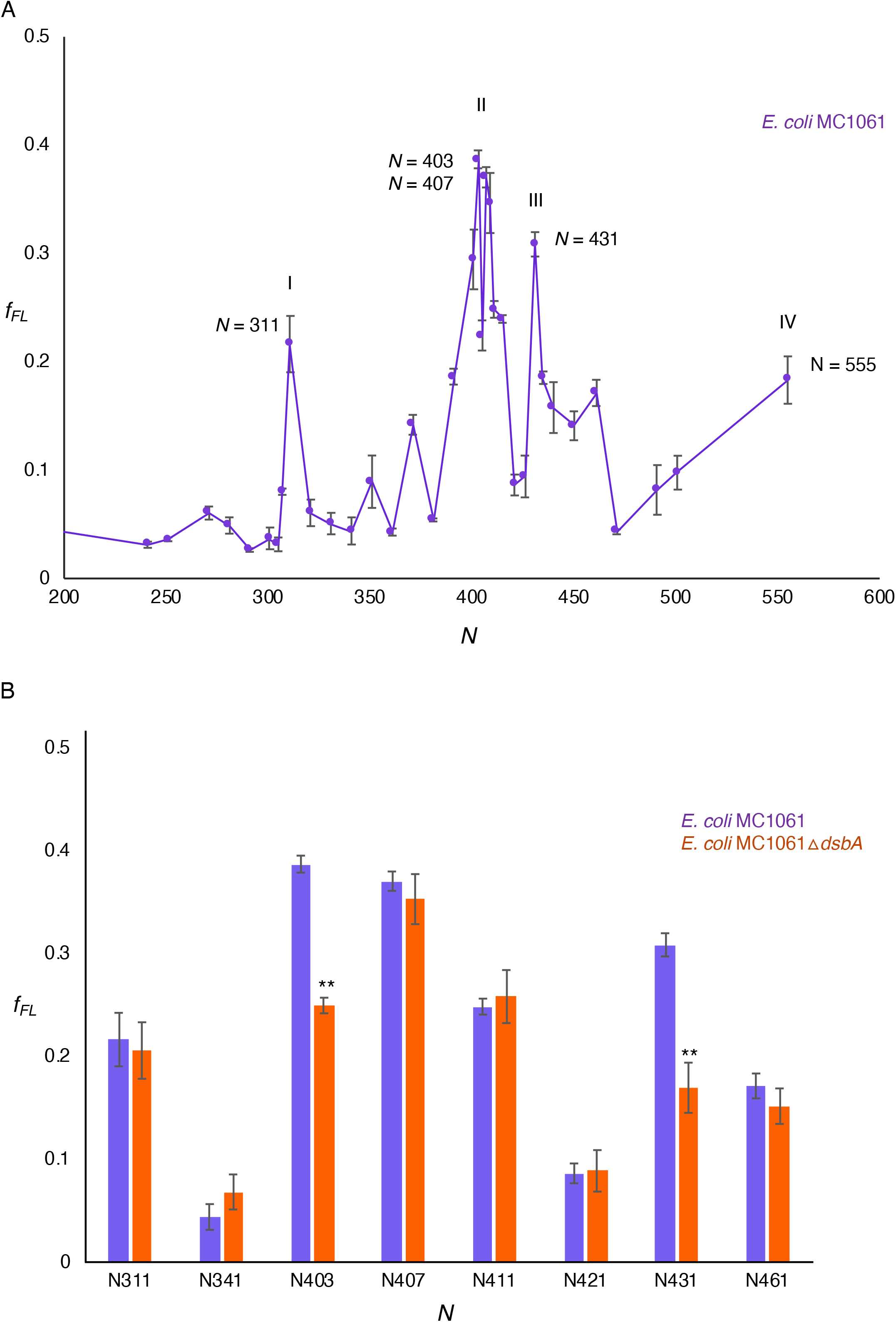
(A) Force profile for PhoA (*N* = 141 to *N* = 555) obtained after pulse-labeling in *E. coli* MC1061. Four maxima (I, II, III, IV) can be seen. The corresponding lengths are indicated in the figure. (B) Comparison of *f*_*FL*_ of selected constructs pulse-labeled in *E. coli* MC1061 (purple) and *E. coli* MC1061 *ΔdsbA* (red). Significant reductions in *f*_*FL*_ (2-sided t-test, *p* < 0.01, indicated as **) were seen for constructs *N* = 403 and 431. Error bars indicate SEM values calculated from independent triplicates of all constructs.

The tightly spaced peaks II and III at *N* ≈ 400-435 residues are particularly interesting, since they correspond well to the respective lengths of PhoA nascent chains (~39 kDa and ~44 kDa, not including the signal peptide) when the cotranslational formation of the C190-C200 and C308-C358 disulfide bonds (catalyzed by DsbA) can first be detected ^43^. Given that an extended polypeptide of ~50 residues length should be able to span the ~160 Å distance between the PTC and the periplasmic exit from the SecYEG translocation channel ^57^, C308 should emerge into the periplasm at *N* ≈ 360 residues, and C358 at *N* ≈ 410 residues. This suggested to us that peaks II and III may reflect the formation of disulfide-stabilized folding intermediates in the periplasm. This is also supported by Fig. 1D, and by *in vivo* studies that have shown that both the C190-C200 and the C308-C358 disulfide bonds are needed for mature PhoA to reach full stability ^58^.

To test this hypothesis, we first recorded *f*_*FL*_ values in and around peaks I, II, and III in the MC1061Δ*dsbA* strain, Fig. 2B. Significant reductions in *f*_*FL*_ (*p* < 0.01, 2-sided t-test) were seen for constructs *N* = 403 and *N* = 431. We further mutated the four Cys residues involved in disulfide-bond formation (C190, C200, C308, C358) to Ala, either individually or in combination, in constructs *N* = 311 (peak I), 403 and 407 (peak II), and 431 (peak III), Fig. 3A-D. In the *N* = 311 and *N* = 403 construct, none of the mutations gave rise to a significant reduction in *f*_*FL*_. Mutations C190A and C200A both significantly reduced *f*_*FL*_ in the *N* = 407 construct. Finally, in the *N* = 431 construct, mutations C200A, C308A, and C358A all strongly reduced *f*_*FL*_, as did the triple mutation C190A+C308A+C358A but not the double mutations C190A+C200A and C308A+C358A.

**Figure 3.**
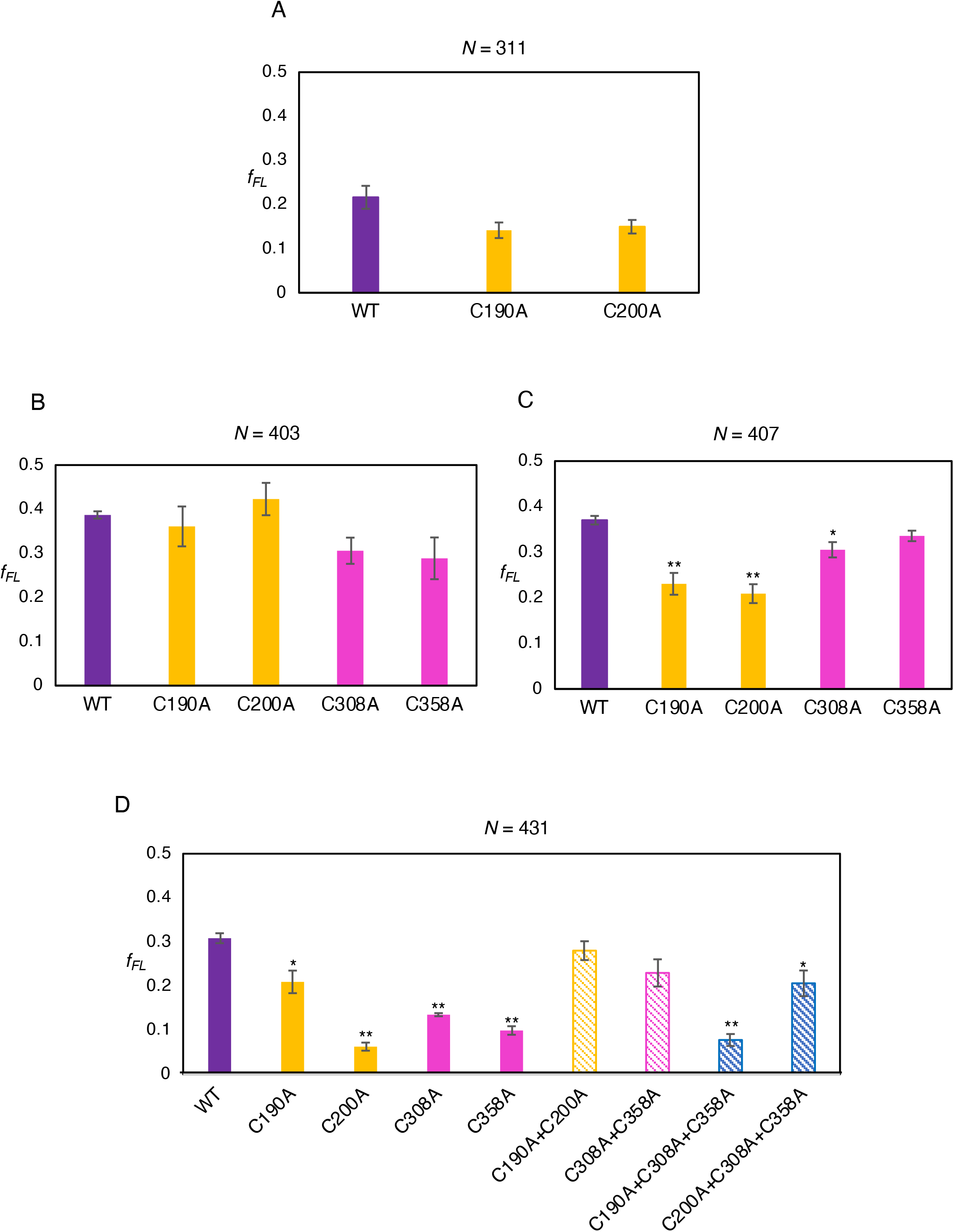
*f*_*FL*_ values for the indicated PhoA Cys→Ala mutations obtained after pulse-labeling in *E. coli* MC1061. (A) For length *N* = 311, no significant drop in *f*_*FL*_ is observed for the individual mutants C190A and C200A (orange bars) compared to the wild-type PhoA truncate *N* = 311 (purple bar). C190A and C200A form the first pair of disulfide bonds in PhoA and are exposed to the periplasm at this length. (B) For length *N* = 403, no significant effects are seen for the individual mutants C190A and C200A (orange bars) or C308A and C358A (that form the second disulfide bond; magenta bars) compared to the wild-type PhoA truncate. C190A, C200A and C308A should be in the periplasm at this length. (C) At *N* = 407, a significant reduction (2-sided t-test, *p* < 0.01, indicated as **) was observed for C190A and C200A (orange bars) compared to the wild-type PhoA truncate (purple bar). No significant reduction was seen for C308A and C358A (magenta bars). (D) At *N* = 431, all four Cys residues are exposed to the periplasm, and the consequences of mutating them to Ala result in significant reductions (*p* < 0.01, indicated as **) for C200A (orange solid bar), C308A, and C358A (magenta solid bars). No significant reduction was seen for the double mutations C190A-C200A (orange striped bar) or C308A-C358A (magenta striped bar). A significant reduction (*p* < 0.01, indicated as **) was observed for the triple mutant C190A-C308A-C358A, (blue striped bar). Error bars indicate SEM values calculated from independent triplicates of all constructs.

The data for peak II (*N* = 407) suggest the existence of an early folding intermediate encompassing residues 1 to ~335 in mature PhoA, Fig. 4A, stabilized by the C190-C200 disulfide bond. At this length, C358 has not yet emerged into the periplasm, making the intermediate sensitive to mutations in the first disulfide (C190, C200) but not in the second (C308, C358). Likewise, the data for peak III (*N* = 431) suggest the existence of a second intermediate encompassing residues 1 to ~360, Fig. 4B, in which all four Cys residues have emerged into the periplasm. In this case, the single Cys→Ala mutations destabilize the folded state both directly and also indirectly through the formation of mis-paired disulfides between the three remaining Cys residues^43,58^. The double mutations C190+C200 and C308+C358 are less detrimental as they leave one native disulfide intact and do not lead to mis-pairing, while the triple mutations prevent both native disulfides from forming. Whether peak I also reflects the formation of a folding intermediate remains unclear, but in any case disulfide bond formation appears not to be involved (Fig. 2B and 3A).

**Figure 4.**
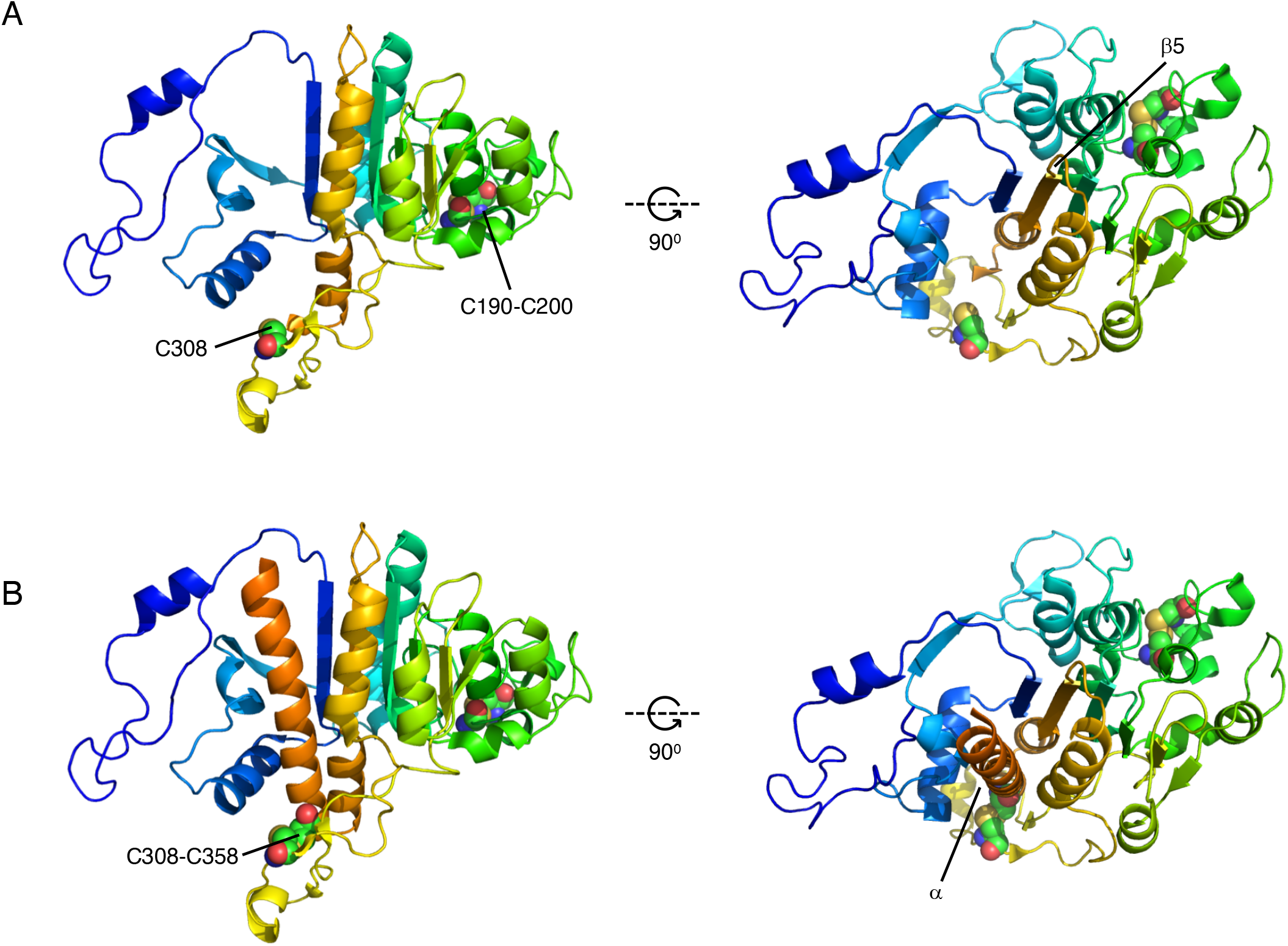
Proposed disulfide-stabilized, cotranslational folding intermediates. (A) Mature PhoA residues 1-335 (peak II). The intermediate represents a state where strand β6 has inserted between strands β1 and β2, completing the β1-β6-β2-β3-β5-β4 portion of the central βsheet. (B) Mature PhoA residues 1-360 (peak III). The intermediate represents a state where the α-helix (in orange) following β6 has packed on top of the central β-sheet. Cys residues are indicated in spacefill and numbered according to UniProt ID P00634, *i.e*., counting from the N-terminus of the signal peptide (which is 22 residues away from the N-terminus of the mature protein). Prepared using the PyMol molecular graphics software (https://pymol.org).

We conclude that FPA can detect long-range forces on the nascent chain generated by cotranslational protein folding in the periplasm, and that PhoA folds cotranslationally via at least two disulfide-stabilized intermediate states. It will be interesting to further investigate the folding of periplasmic proteins with non-consecutive disulfide bonds, and to dissect the role of periplasmic chaperones in cotranslational folding.

## Materials and Methods

### Reagents, chemicals, and primers

All chemicals and reagents were purchased from Merck Sigma Aldrich; primers for PCR and DNA sequencing were purchased from Eurofins Genomics; gene fragments, Bis-Tris gels, Phusion DNA polymerase, GeneJet plasmid isolation kits, GeneJet PCR purification kits, and GeneJet Gel extraction kits were purchased from Thermo Fisher Scientific; [^35^S] Methionine was purchased from Perkin Elmer; Protein-G agarose beads were manufactured by Roche and purchased from Sigma Aldrich; the mouse monoclonal antibody against the HA antigen was purchased from Covance.

### Design and engineering of full-length PhoA fused to the SecM AP

The gene sequence encoding full-length PhoA-GSGS-HA-SecM was designed *in silico* and ordered from GeneArt, Thermo Fisher Scientific.

The construct was designed as follows: the sequence of the *phoA* gene from *E. coli* K12 MG1655 was obtained from the *EcoCyc* database^59^, and was engineered upstream of sequences encoding a four amino-acid long GSGS linker, an HA tag, the 17 amino acid long *E. coli* SecM AP, and a C-terminal tail consisting of 77 residues derived from the sequence of LacZ.

The obtained gene fragment was engineered by Gibson Assembly^60^ into a previously described plasmid that harbors an arabinose-inducible promoter and an ampicillin resistance gene for selection^49^, and sequenced for verification.

### Engineering a non-cleavable PhoA signal and generation of truncates

The non-cleavable (NC) PhoA signal peptide (MKQSTIALALLPLLFTPVTKPR) was designed based on predictions using SignalP 5.0 ^61^ and PhoA mutants with non-cleavable signal peptides ^56^. The sequence used consists of a single amino acid substitution (A to P) at residue 21 in PhoA (underlined above). This single substitution was introduced into the gene sequence encoding PhoA-GSGS-HA-SecM using partially overlapping PCR and the following pair of primers:

forward 5’-GTGACAAAACCACGGACACCAGAAATGCC-3’ and,
reverse 5’-TTCTGGTGTCCGTGGTTTTGTCACAGGGGTAAA-3’.

The resulting parental construct - NCPhoA-GSGS-HA-SecM - was used to engineer constructs that were truncated every ten amino acid residues from the the C-terminus of PhoA (primer list and protein sequences provided as Supplementary Information).

The resulting PCR products were subjected to *Dpn1* digestion and transformed into DH5α cells, thereafter the plasmids were isolated and sequences verified (Eurofins Genomics).

Plasmids bearing the correct sequences were transformed into *E. coli* MC1061 competent cells for *in vivo* expression and pulse labelling.

### Engineering the E. coli MC1061ΔdsbA strain

The MC1061Δ*dsbA* strain was engineered using the λ-Red recombineering-based approach developed by Datsenko and Wanner^62^. In short, a kanamycin cassette with flanking regions homologous to the up- and downstream regions of the *dsbA* gene was generated by PCR using the pKD13 plasmid as a template and the dsbA-FRT-fw and dsbA-FRT-rv primers. The PCR-reaction was treated with *Dpn*I (NEB CutSmart) to get rid of the template DNA. The PCR reaction was loaded on an agarose gel to verify the molecular weight of the generated PCR product and it was subsequently purified using the GeneJET PCR Purification kit (Thermo Scientific). The purified PCR product was electroporated into *E. coli* MC1061 cells harboring pKD46 that were cultured in standard Lysogeny Broth (LB) in the presence of 0.2% arabinose at 30° C. After verifying the correct insertion of the kanamycin cassette into the *dsbA* gene using colony PCR and the primers dsbA-up and dsbA-down, *dsbA*::Km^R^ was transferred from MC1061Δ*dsbA*::Km^R^/pKD46 to wild-type MC1061 by means of P1-transduction ^63^ to generate MC1061Δ*dsbA*::Km^R^

dsbA-FRT-rv: AGAACCCCCTTTGCAATTAACACCTATGTATTAATCGGAGAGAGTAGATCTGTAGGCTGGAGCTGCTTCG
dsbA-FRT-fw: TAATAAAAAAAGCCCGTGAATATTCACGGGCTTTATGTAATTTACATTGAAATTCCGGGGATCCGTCGACC
dsbA-up: TACGGCTAACGCAACAATAACACC
dsbA-dw: CATTCCTGAAAGCGACAGATGAG

### In vivo expression and pulse labelling

*E. coli* MC1061 or MC1061Δ*dsbA* harboring arabinose-inducible plasmids with the PhoA constructs of interest were cultured for 14 hours in 2 ml LB supplemented with 100 μg/ml Ampicillin at 37° C and shaking at 200 rpm in a New Brunswick Scientific Innova 44 incubator shaker.

After approx. 14 hours, the cultures were back-diluted into 2 ml LB (supplemented with 100 μg/ml Ampicillin) to an A_600_ of 0.1, and the fresh suspensions were cultured at 37° C with shaking at 200 rpm till they reached an A600 of 0.4 (measured using a Shimadzu UV-1300 spectrophotometer). The cultures were chilled on ice for 5 minutes, transferred into 2 ml reaction tubes (Sarstedt), and cells were isolated by centrifugation at 5000×*g* for 8 minutes in a cooled table-top centrifuge. The supernatant was discarded rapidly, and the cell pellets were washed in M9 minimal medium (supplemented with 1 μg/ml of 19 amino acids excluding Methionine, 100 μg/ml Thiamine, 0.1 mM CaCl_2_, 2 mM MgSO_4_, 0.4% w/w fructose, and 100 μg/ml Ampicillin), by gentle resuspension.

The suspensions were centrifuged once again to obtain cell pellets, which were subsequently resuspended in M9 medium at 37° C. These were cultured for one hour at 37° C and shaking at 200 rpm.

The cell cultures were then transferred into 2 ml reaction tubes and placed in slots in a table-top thermomixer (Eppendorf) set to 37° C and shaking at 700 rpm for five minutes prior to pulse-labelling. Expression of the gene of interest was induced by 0.2% Arabinose for five minutes, and cells were pulse-labelled with 4 μCi [^35^S] Methionine for two minutes. The reaction was aborted by treating the cells with ice-cold Trichloroacetic acid (TCA) at a final conc. of 10%, and incubating at −20° C for 20 min. The precipitated samples were centrifuged for 10 minutes at 20,000x*g* at 4° C in a table-top centrifuge.

The resulting pellets were washed with ice-cold acetone (to neutralize the TCA and solvate lipids) and centrifuged at 20,000×*g* for 10 minutes in a table-top micro-centrifuge (Eppendorf). The obtained pellets were air-dried to rid them of residual acetone and solubilized with 120 μl Tris-SDS (10 mM Tris-HCl pH 7.5 and 2% SDS) at 56° C for 10 min and shaking at 1000 rpm in a table-top thermomixer (Eppendorf). The resulting suspension was centrifuged 20,000×*g* for 10 minutes at room temperature in a table-top micro-centrifuge (Eppendorf), and 100 μl of the supernatant was subjected to immune precipitation using the anti-HA antiserum.

The obtained samples were separated by SDS-PAGE and visualized by autoradiography on a FujiFilm Scanner FLA-9000.

### Quantification of radioactively labelled proteins

The *FL* and *A* protein bands on the gel visualized after autoradiography were quantified using MultiGauge (Fujifilm) from which one-dimensional intensity profile of each gel lane was extracted. The band intensities were fit to Gaussian distributions using EasyQuant ^54^. The sum of the arrested and full-length band intensities was calculated, and this was used to estimate the fraction of full-length protein for each construct. Independent triplicates were run for all constructs, and averages and SEM values were calculated.

## Supporting information

Amino acid and primer sequences

## Acknowledgements

This work was supported by grants from the Knut and Alice Wallenberg Foundation (2017.0323), the Novo Nordisk Fund (NNF18OC0032828), and the Swedish Research Council (621-2014-3713) to GvH, and the Swedish Research Council (2019-04143) and the Novo Nordisk Fund (NNF19OC0057673) to JWdG.

