## Supplementary material for "Cotranslational folding of alkaline phosphatase in the periplasm of *Escherichia coli*": Amino acid and primer sequences

**Sequences of the NC PhoA constructs**

**Key:** PhoA NC Signal Sequence – PhoA mature domain – SGSG – Linker extension – HA tag – SecM AP – C-terminal tail

| Name | Sequence |
|---|---|
| N= 555 | MKQSTIALALLPLLFTPVTKPRTPEMPVLENRAAQGDITAPGGARRLTG<br>DQTAALRDSLSDKPAKNIILLIGDGMGDSEITAARNYAEGAGGFFKGID<br>ALPLTGQYTHYALNKKTGKPDYVTDSAASATAWSTGVKTYNGALGVDIH<br>EKDHPTILEMAKAAGLATGNVSTAELQDATPAALVAHVTSRKCYGPSAT<br>SEKCPGNALEKGGKGSITEQLLNARADVTLGGGAKTFAETATAGEWQGK<br>TLREQAQARGYQLVSDAASLNSVTEANQQKPLLGLFADGNMPVRWLGPK<br>ATYHGNIDKPAVTCTPNPQRNDSVPTLAQMTDKAIELLSKNEKGFFLQV<br>EGASIDKQDHAANPCGQIGETVDLDEAVQRALEFAKKEGNTLVIVTADH<br>AHASQIVAPDTKAPGLTQALNTKDGAVMVMSYGNSEEDSQEHTGSQLRI<br>AAYGPHAANVVGLTDQTDLFYTMKAALGLKSGSGSGSGYPYDVPDYAMY<br>PYDVPDYAMYPYDVPDYAMYPYDVPDYAMYPYDVPDYAMYPYDVPDYAM<br>YFSTPVWISQAQGIRAGPGSSDKQEGEWPTGLRLSRIGGIHSLAVVLQR<br>RDWENPGVTQLNRLAAHPPFASWRNSEEARTDRPSQQLRSLNGEWR* |
| N= 501 | MKQSTIALALLPLLFTPVTKPRMKQSTIALALLPLLFTPVTKPRTPEMP<br>VLENRAAQGDITAPGGARRLTGDQTAALRDSLSDKPAKNIILLIGDGMG<br>DSEITAARNYAEGAGGFFKGIDALPLTGQYTHYALNKKTGKPDYVTDSA<br>ASATAWSTGVKTYNGALGVDIHEKDHPTILEMAKAAGLATGNVSTAELQ<br>DATPAALVAHVTSRKCYGPSATSEKCPGNALEKGGKGSITEQLLNARAD<br>VTLGGGAKTFAETATAGEWQGKTLREQAQARGYQLVSDAASLNSVTEAN<br>QQKPLLGLFADGNMPVRWLGPKATYHGNIDKPAVTCTPNPQRNDSVPTL<br>AQMTDKAIELLSKNEKGFFLQVEGASIDKQDHAANPCGQIGETVDLDEA<br>VQRALEFAKKEGNTLVIVTADHAHASQIVAPDTKAPGLTQALNTKDGAV<br>MVMSYGNSEEDSQEHTGSQLRIAAYGPHAANVVGLTDQTDLFYTMKAAL<br>GLKSGSGYPYDVPDYAFSTPVWISQAQGIRAGPGSSDKQEGEWPTGLRL<br>SRIGGIHSLAVVLQRRDWENPGVTQLNRLAAHPPFASWRNSEEARTDRP<br>SQQLRSLNGEWR* |
| N= 491 | MKQSTIALALLPLLFTPVTKPRMKQSTIALALLPLLFTPVTKPRTPEMP<br>VLENRAAQGDITAPGGARRLTGDQTAALRDSLSDKPAKNIILLIGDGMG<br>DSEITAARNYAEGAGGFFKGIDALPLTGQYTHYALNKKTGKPDYVTDSA<br>ASATAWSTGVKTYNGALGVDIHEKDHPTILEMAKAAGLATGNVSTAELQ<br>DATPAALVAHVTSRKCYGPSATSEKCPGNALEKGGKGSITEQLLNARAD<br>VTLGGGAKTFAETATAGEWQGKTLREQAQARGYQLVSDAASLNSVTEAN<br>QQKPLLGLFADGNMPVRWLGPKATYHGNIDKPAVTCTPNPQRNDSVPTL<br>AQMTDKAIELLSKNEKGFFLQVEGASIDKQDHAANPCGQIGETVDLDEA<br>VQRALEFAKKEGNTLVIVTADHAHASQIVAPDTKAPGLTQALNTKDGAV<br>MVMSYGNSEEDSQEHTGSQLRIAAYGPHAANVVGLTDQTDLFSGSGYPY<br>DVPDYAFSTPVWISQAQGIRAGPGSSDKQEGEWPTGLRLSRIGGIHSLA<br>VVLQRRDWENPGVTQLNRLAAHPPFASWRNSEEARTDRPSQQLRSLNGE<br>WR* |
| N= 471 | MKQSTIALALLPLLFTPVTKPRMKQSTIALALLPLLFTPVTKPRTPEMP<br>VLENRAAQGDITAPGGARRLTGDQTAALRDSLSDKPAKNIILLIGDGMG<br>DSEITAARNYAEGAGGFFKGIDALPLTGQYTHYALNKKTGKPDYVTDSA<br>ASATAWSTGVKTYNGALGVDIHEKDHPTILEMAKAAGLATGNVSTAELQ<br>DATPAALVAHVTSRKCYGPSATSEKCPGNALEKGGKGSITEQLLNARAD<br>VTLGGGAKTFAETATAGEWQGKTLREQAQARGYQLVSDAASLNSVTEAN<br>QQKPLLGLFADGNMPVRWLGPKATYHGNIDKPAVTCTPNPQRNDSVPTL<br>AQMTDKAIELLSKNEKGFFLQVEGASIDKQDHAANPCGQIGETVDLDEA |

| | |
|---|---|
| | VQRALEFAKKEGNTLVIVTADHAHASQIVAPDTKAPGLTQALNTKDGAV<br>MVMSYGNSEEDSQEHTGSQLRISGSGYPYDVPDYAFSTPVWISQAQGIR<br>AGPGSSDKQEGEWPTGLRLSRIGGIHSLAVVLQRRDWENPGVTQLNRLA<br>AHPPFASWRNSEEARTDRPSQQLRSLNGEWR* |
| N= 461 | MKQSTIALALLPLLFTPVTKPRMKQSTIALALLPLLFTPVTKPRTPEMP<br>VLENRAAQGDITAPGGARRLTGDQTAALRDSLSDKPAKNIILLIGDGMG<br>DSEITAARNYAEGAGGFFKGIDALPLTGQYTHYALNKKTGKPDYVTDSA<br>ASATAWSTGVKTYNGALGVDIHEKDHPTILEMAKAAGLATGNVSTAELQ<br>DATPAALVAHVTSRKCYGPSATSEKCPGNALEKGGKGSITEQLLNARAD<br>VTLGGGAKTFAETATAGEWQGKTLREQAQARGYQLVSDAASLNSVTEAN<br>QQKPLLGLFADGNMPVRWLGPKATYHGNIDKPAVTCTPNPQRNDSVPTL<br>AQMTDKAIELLSKNEKGFFLQVEGASIDKQDHAANPCGQIGETVDLDEA<br>VQRALEFAKKEGNTLVIVTADHAHASQIVAPDTKAPGLTQALNTKDGAV<br>MVMSYGNSEEDSSGSGYPYDVPDYAFSTPVWISQAQGIRAGPGSSDKQE<br>GEWPTGLRLSRIGGIHSLAVVLQRRDWENPGVTQLNRLAAHPPFASWRN<br>SEEARTDRPSQQLRSLNGEWR* |
| N= 450 | MKQSTIALALLPLLFTPVTKPRMKQSTIALALLPLLFTPVTKPRTPEMP<br>VLENRAAQGDITAPGGARRLTGDQTAALRDSLSDKPAKNIILLIGDGMG<br>DSEITAARNYAEGAGGFFKGIDALPLTGQYTHYALNKKTGKPDYVTDSA<br>ASATAWSTGVKTYNGALGVDIHEKDHPTILEMAKAAGLATGNVSTAELQ<br>DATPAALVAHVTSRKCYGPSATSEKCPGNALEKGGKGSITEQLLNARAD<br>VTLGGGAKTFAETATAGEWQGKTLREQAQARGYQLVSDAASLNSVTEAN<br>QQKPLLGLFADGNMPVRWLGPKATYHGNIDKPAVTCTPNPQRNDSVPTL<br>AQMTDKAIELLSKNEKGFFLQVEGASIDKQDHAANPCGQIGETVDLDEA<br>VQRALEFAKKEGNTLVIVTADHAHASQIVAPDTKAPGLTQALNTKDGAV<br>MSGSGYPYDVPDYAFSTPVWISQAQGIRAGPGSSDKQEGEWPTGLRLSR<br>IGGIHSLAVVLQRRDWENPGVTQLNRLAAHPPFASWRNSEEARTDRPSQ<br>QLRSLNGEWR* |
| N= 440 | MKQSTIALALLPLLFTPVTKPRMKQSTIALALLPLLFTPVTKPRTPEMP<br>VLENRAAQGDITAPGGARRLTGDQTAALRDSLSDKPAKNIILLIGDGMG<br>DSEITAARNYAEGAGGFFKGIDALPLTGQYTHYALNKKTGKPDYVTDSA<br>ASATAWSTGVKTYNGALGVDIHEKDHPTILEMAKAAGLATGNVSTAELQ<br>DATPAALVAHVTSRKCYGPSATSEKCPGNALEKGGKGSITEQLLNARAD<br>VTLGGGAKTFAETATAGEWQGKTLREQAQARGYQLVSDAASLNSVTEAN<br>QQKPLLGLFADGNMPVRWLGPKATYHGNIDKPAVTCTPNPQRNDSVPTL<br>AQMTDKAIELLSKNEKGFFLQVEGASIDKQDHAANPCGQIGETVDLDEA<br>VQRALEFAKKEGNTLVIVTADHAHASQIVAPDTKAPGLTQSGSGYPYDV<br>PDYAFSTPVWISQAQGIRAGPGSSDKQEGEWPTGLRLSRIGGIHSLAVV<br>LQRRDWENPGVTQLNRLAAHPPFASWRNSEEARTDRPSQQLRSLNGEWR* |
| N= 435 | MKQSTIALALLPLLFTPVTKPRMKQSTIALALLPLLFTPVTKPRTPEMP<br>VLENRAAQGDITAPGGARRLTGDQTAALRDSLSDKPAKNIILLIGDGMG<br>DSEITAARNYAEGAGGFFKGIDALPLTGQYTHYALNKKTGKPDYVTDSA<br>ASATAWSTGVKTYNGALGVDIHEKDHPTILEMAKAAGLATGNVSTAELQ<br>DATPAALVAHVTSRKCYGPSATSEKCPGNALEKGGKGSITEQLLNARAD<br>VTLGGGAKTFAETATAGEWQGKTLREQAQARGYQLVSDAASLNSVTEAN<br>QQKPLLGLFADGNMPVRWLGPKATYHGNIDKPAVTCTPNPQRNDSVPTL<br>AQMTDKAIELLSKNEKGFFLQVEGASIDKQDHAANPCGQIGETVDLDEA<br>VQRALEFAKKEGNTLVIVTADHAHASQIVAPDTKASGSGYPYDVPDYAF<br>STPVWISQAQGIRAGPGSSDKQEGEWPTGLRLSRIGGIHSLAVVLQRRD<br>WENPGVTQLNRLAAHPPFASWRNSEEARTDRPSQQLRSLNGEWR* |
| N= 431 | MKQSTIALALLPLLFTPVTKPRMKQSTIALALLPLLFTPVTKPRTPEMP<br>VLENRAAQGDITAPGGARRLTGDQTAALRDSLSDKPAKNIILLIGDGMG |

| | |
|---|---|
| | DSEITAARNYAEGAGGFFKGIDALPLTGQYTHYALNKKTGKPDYVTDSA<br>ASATAWSTGVKTYNGALGVDIHEKDHPTILEMAKAAGLATGNVSTAELQ<br>DATPAALVAHVTSRKCYGPSATSEKCPGNALEKGGKGSITEQLLNARAD<br>VTLGGGAKTFAETATAGEWQGKTLREQAQARGYQLVSDAASLNSVTEAN<br>QQKPLLGLFADGNMPVRWLGPKATYHGNIDKPAVTCTPNPQRNDSVPTL<br>AQMTDKAIELLSKNEKGFFLQVEGASIDKQDHAANPCGQIGETVDLDEA<br>VQRALEFAKKEGNTLVIVTADHAHASQIVAPSGSGYPYDVPDYAFSTPV<br>WISQAQGIRAGPGSSDKQEGEWPTGLRLSRIGGIHSLAVVLQRRDWENP<br>GVTQLNRLAAHPPFASWRNSEEARTDRPSQQLRSLNGEWR* |
| N= 426 | MKQSTIALALLPLLFTPVTKPRMKQSTIALALLPLLFTPVTKPRTPEMP<br>VLENRAAQGDITAPGGARRLTGDQTAALRDSLSDKPAKNIILLIGDGMG<br>DSEITAARNYAEGAGGFFKGIDALPLTGQYTHYALNKKTGKPDYVTDSA<br>ASATAWSTGVKTYNGALGVDIHEKDHPTILEMAKAAGLATGNVSTAELQ<br>DATPAALVAHVTSRKCYGPSATSEKCPGNALEKGGKGSITEQLLNARAD<br>VTLGGGAKTFAETATAGEWQGKTLREQAQARGYQLVSDAASLNSVTEAN<br>QQKPLLGLFADGNMPVRWLGPKATYHGNIDKPAVTCTPNPQRNDSVPTL<br>AQMTDKAIELLSKNEKGFFLQVEGASIDKQDHAANPCGQIGETVDLDEA<br>VQRALEFAKKEGNTLVIVTADHAHASSGSGYPYDVPDYAFSTPVWISQA<br>QGIRAGPGSSDKQEGEWPTGLRLSRIGGIHSLAVVLQRRDWENPGVTQL<br>NRLAAHPPFASWRNSEEARTDRPSQQLRSLNGEWR* |
| N= 421 | MKQSTIALALLPLLFTPVTKPRMKQSTIALALLPLLFTPVTKPRTPEMP<br>VLENRAAQGDITAPGGARRLTGDQTAALRDSLSDKPAKNIILLIGDGMG<br>DSEITAARNYAEGAGGFFKGIDALPLTGQYTHYALNKKTGKPDYVTDSA<br>ASATAWSTGVKTYNGALGVDIHEKDHPTILEMAKAAGLATGNVSTAELQ<br>DATPAALVAHVTSRKCYGPSATSEKCPGNALEKGGKGSITEQLLNARAD<br>VTLGGGAKTFAETATAGEWQGKTLREQAQARGYQLVSDAASLNSVTEAN<br>QQKPLLGLFADGNMPVRWLGPKATYHGNIDKPAVTCTPNPQRNDSVPTL<br>AQMTDKAIELLSKNEKGFFLQVEGASIDKQDHAANPCGQIGETVDLDEA<br>VQRALEFAKKEGNTLVIVTADSGSGYPYDVPDYAFSTPVWISQAQGIRA<br>GPGSSDKQEGEWPTGLRLSRIGGIHSLAVVLQRRDWENPGVTQLNRLAA<br>HPPFASWRNSEEARTDRPSQQLRSLNGEWR* |
| N= 415 | MKQSTIALALLPLLFTPVTKPRMKQSTIALALLPLLFTPVTKPRTPEMP<br>VLENRAAQGDITAPGGARRLTGDQTAALRDSLSDKPAKNIILLIGDGMG<br>DSEITAARNYAEGAGGFFKGIDALPLTGQYTHYALNKKTGKPDYVTDSA<br>ASATAWSTGVKTYNGALGVDIHEKDHPTILEMAKAAGLATGNVSTAELQ<br>DATPAALVAHVTSRKCYGPSATSEKCPGNALEKGGKGSITEQLLNARAD<br>VTLGGGAKTFAETATAGEWQGKTLREQAQARGYQLVSDAASLNSVTEAN<br>QQKPLLGLFADGNMPVRWLGPKATYHGNIDKPAVTCTPNPQRNDSVPTL<br>AQMTDKAIELLSKNEKGFFLQVEGASIDKQDHAANPCGQIGETVDLDEA<br>VQRALEFAKKEGNTLSGSGYPYDVPDYAFSTPVWISQAQGIRAGPGSSD<br>KQEGEWPTGLRLSRIGGIHSLAVVLQRRDWENPGVTQLNRLAAHPPFAS<br>WRNSEEARTDRPSQQLRSLNGEWR* |
| N= 411 | MKQSTIALALLPLLFTPVTKPRMKQSTIALALLPLLFTPVTKPRTPEMP<br>VLENRAAQGDITAPGGARRLTGDQTAALRDSLSDKPAKNIILLIGDGMG<br>DSEITAARNYAEGAGGFFKGIDALPLTGQYTHYALNKKTGKPDYVTDSA<br>ASATAWSTGVKTYNGALGVDIHEKDHPTILEMAKAAGLATGNVSTAELQ<br>DATPAALVAHVTSRKCYGPSATSEKCPGNALEKGGKGSITEQLLNARAD<br>VTLGGGAKTFAETATAGEWQGKTLREQAQARGYQLVSDAASLNSVTEAN<br>QQKPLLGLFADGNMPVRWLGPKATYHGNIDKPAVTCTPNPQRNDSVPTL<br>AQMTDKAIELLSKNEKGFFLQVEGASIDKQDHAANPCGQIGETVDLDEA<br>VQRALEFAKKESGSGYPYDVPDYAFSTPVWISQAQGIRAGPGSSDKQEG<br>EWPTGLRLSRIGGIHSLAVVLQRRDWENPGVTQLNRLAAHPPFASWRNS |

| | |
|---|---|
| | EEARTDRPSQQLRSLNGEWR* |
| N= 409 | MKQSTIALALLPLLFTPVTKPRMKQSTIALALLPLLFTPVTKPRTPEMP<br>VLENRAAQGDITAPGGARRLTGDQTAALRDSLSDKPAKNIILLIGDGMG<br>DSEITAARNYAEGAGGFFKGIDALPLTGQYTHYALNKKTGKPDYVTDSA<br>ASATAWSTGVKTYNGALGVDIHEKDHPTILEMAKAAGLATGNVSTAELQ<br>DATPAALVAHVTSRKCYGPSATSEKCPGNALEKGGKGSITEQLLNARAD<br>VTLGGGAKTFAETATAGEWQGKTLREQAQARGYQLVSDAASLNSVTEAN<br>QQKPLLGLFADGNMPVRWLGPKATYHGNIDKPAVTCTPNPQRNDSVPTL<br>AQMTDKAIELLSKNEKGFFLQVEGASIDKQDHAANPCGQIGETVDLDEA<br>VQRALEFAKSGSGYPYDVPDYAFSTPVWISQAQGIRAGPGSSDKQEGEW<br>PTGLRLSRIGGIHSLAVVLQRRDWENPGVTQLNRLAAHPPFASWRNS<br>EEARTDRPSQQLRSLNGEWR* |
| N= 407 | MKQSTIALALLPLLFTPVTKPRMKQSTIALALLPLLFTPVTKPRTPEMP<br>VLENRAAQGDITAPGGARRLTGDQTAALRDSLSDKPAKNIILLIGDGMG<br>DSEITAARNYAEGAGGFFKGIDALPLTGQYTHYALNKKTGKPDYVTDSA<br>ASATAWSTGVKTYNGALGVDIHEKDHPTILEMAKAAGLATGNVSTAELQ<br>DATPAALVAHVTSRKCYGPSATSEKCPGNALEKGGKGSITEQLLNARAD<br>VTLGGGAKTFAETATAGEWQGKTLREQAQARGYQLVSDAASLNSVTEAN<br>QQKPLLGLFADGNMPVRWLGPKATYHGNIDKPAVTCTPNPQRNDSVPTL<br>AQMTDKAIELLSKNEKGFFLQVEGASIDKQDHAANPCGQIGETVDLDEA<br>VQRALEFSGSGYPYDVPDYAFSTPVWISQAQGIRAGPGSSDKQEGEWPT<br>GLRLSRIGGIHSLAVVLQRRDWENPGVTQLNRLAAHPPFASWRNSEEAR<br>TDRPSQQLRSLNGEWR* |
| N= 405 | MKQSTIALALLPLLFTPVTKPRMKQSTIALALLPLLFTPVTKPRTPEMP<br>VLENRAAQGDITAPGGARRLTGDQTAALRDSLSDKPAKNIILLIGDGMG<br>DSEITAARNYAEGAGGFFKGIDALPLTGQYTHYALNKKTGKPDYVTDSA<br>ASATAWSTGVKTYNGALGVDIHEKDHPTILEMAKAAGLATGNVSTAELQ<br>DATPAALVAHVTSRKCYGPSATSEKCPGNALEKGGKGSITEQLLNARAD<br>VTLGGGAKTFAETATAGEWQGKTLREQAQARGYQLVSDAASLNSVTEAN<br>QQKPLLGLFADGNMPVRWLGPKATYHGNIDKPAVTCTPNPQRNDSVPTL<br>AQMTDKAIELLSKNEKGFFLQVEGASIDKQDHAANPCGQIGETVDLDEA<br>VQRALSGSGYPYDVPDYAFSTPVWISQAQGIRAGPGSSDKQEGEWPTGL<br>RLSRIGGIHSLAVVLQRRDWENPGVTQLNRLAAHPPFASWRNSEEARTD<br>RPSQQLRSLNGEWR* |
| N= 403 | MKQSTIALALLPLLFTPVTKPRMKQSTIALALLPLLFTPVTKPRTPEMP<br>VLENRAAQGDITAPGGARRLTGDQTAALRDSLSDKPAKNIILLIGDGMG<br>DSEITAARNYAEGAGGFFKGIDALPLTGQYTHYALNKKTGKPDYVTDSA<br>ASATAWSTGVKTYNGALGVDIHEKDHPTILEMAKAAGLATGNVSTAELQ<br>DATPAALVAHVTSRKCYGPSATSEKCPGNALEKGGKGSITEQLLNARAD<br>VTLGGGAKTFAETATAGEWQGKTLREQAQARGYQLVSDAASLNSVTEAN<br>QQKPLLGLFADGNMPVRWLGPKATYHGNIDKPAVTCTPNPQRNDSVPTL<br>AQMTDKAIELLSKNEKGFFLQVEGASIDKQDHAANPCGQIGETVDLDEA<br>VQRSGSGYPYDVPDYAFSTPVWISQAQGIRAGPGSSDKQEGEWPTGLRL<br>SRIGGIHSLAVVLQRRDWENPGVTQLNRLAAHPPFASWRNSEEARTDRP<br>SQQLRSLNGEWR* |
| N= 401 | MKQSTIALALLPLLFTPVTKPRMKQSTIALALLPLLFTPVTKPRTPEMP<br>VLENRAAQGDITAPGGARRLTGDQTAALRDSLSDKPAKNIILLIGDGMG<br>DSEITAARNYAEGAGGFFKGIDALPLTGQYTHYALNKKTGKPDYVTDSA<br>ASATAWSTGVKTYNGALGVDIHEKDHPTILEMAKAAGLATGNVSTAELQ<br>DATPAALVAHVTSRKCYGPSATSEKCPGNALEKGGKGSITEQLLNARAD<br>VTLGGGAKTFAETATAGEWQGKTLREQAQARGYQLVSDAASLNSVTEAN<br>QQKPLLGLFADGNMPVRWLGPKATYHGNIDKPAVTCTPNPQRNDSVPTL |

| | |
|---|---|
| | AQMTDKAIELLSKNEKGFFLQVEGASIDKQDHAANPCGQIGETVDLDEA<br>VSGSGYPYDVPDYAFSTPVWISQAQGIRAGPGSSDKQEGEWPTGLRLSR<br>IGGIHSLAVVLQRRDWENPGVTQLNRLAAHPPFASWRNSEEARTDRPS<br>QQLRSLNGEWR* |
| N= 391 | MKQSTIALALLPLLFTPVTKPRMKQSTIALALLPLLFTPVTKPRTPEMP<br>VLENRAAQGDITAPGGARRLTGDQTAALRDSLSDKPAKNIILLIGDGMG<br>DSEITAARNYAEGAGGFFKGIDALPLTGQYTHYALNKKTGKPDYVTDSA<br>ASATAWSTGVKTYNGALGVDIHEKDHPTILEMAKAAGLATGNVSTAELQ<br>DATPAALVAHVTSRKCYGPSATSEKCPGNALEKGGKGSITEQLLNARAD<br>VTLGGGAKTFAETATAGEWQGKTLREQAQARGYQLVSDAASLNSVTEAN<br>QQKPLLGLFADGNMPVRWLGPKATYHGNIDKPAVTCTPNPQRNDSVPTL<br>AQMTDKAIELLSKNEKGFFLQVEGASIDKQDHAANPCGQISGSGYPYDV<br>PDYAFSTPVWISQAQGIRAGPGSSDKQEGEWPTGLRLSRIGGIHSLAVV<br>LQRRDWENPGVTQLNRLAAHPPFASWRNSEEARTDRPSQQLRSLNGEWR<br>* |
| N= 381 | MKQSTIALALLPLLFTPVTKPRMKQSTIALALLPLLFTPVTKPRTPEMP<br>VLENRAAQGDITAPGGARRLTGDQTAALRDSLSDKPAKNIILLIGDGMG<br>DSEITAARNYAEGAGGFFKGIDALPLTGQYTHYALNKKTGKPDYVTDSA<br>ASATAWSTGVKTYNGALGVDIHEKDHPTILEMAKAAGLATGNVSTAELQ<br>DATPAALVAHVTSRKCYGPSATSEKCPGNALEKGGKGSITEQLLNARAD<br>VTLGGGAKTFAETATAGEWQGKTLREQAQARGYQLVSDAASLNSVTEAN<br>QQKPLLGLFADGNMPVRWLGPKATYHGNIDKPAVTCTPNPQRNDSVPTL<br>AQMTDKAIELLSKNEKGFFLQVEGASIDKQSGSGYPYDVPDYAFSTPVW<br>ISQAQGIRAGPGSSDKQEGEWPTGLRLSRIGGIHSLAVVLQRRDWENPG<br>VTQLNRLAAHPPFASWRNSEEARTDRPSQQLRSLNGEWR* |
| N= 371 | MKQSTIALALLPLLFTPVTKPRMKQSTIALALLPLLFTPVTKPRTPEMP<br>VLENRAAQGDITAPGGARRLTGDQTAALRDSLSDKPAKNIILLIGDGMG<br>DSEITAARNYAEGAGGFFKGIDALPLTGQYTHYALNKKTGKPDYVTDSA<br>ASATAWSTGVKTYNGALGVDIHEKDHPTILEMAKAAGLATGNVSTAELQ<br>DATPAALVAHVTSRKCYGPSATSEKCPGNALEKGGKGSITEQLLNARAD<br>VTLGGGAKTFAETATAGEWQGKTLREQAQARGYQLVSDAASLNSVTEAN<br>QQKPLLGLFADGNMPVRWLGPKATYHGNIDKPAVTCTPNPQRNDSVPTL<br>AQMTDKAIELLSKNEKGFFLSGSGYPYDVPDYAFSTPVWISQAQGIRAG<br>PGSSDKQEGEWPTGLRLSRIGGIHSLAVVLQRRDWENPGVTQLNRLAAH<br>PPFASWRNSEEARTDRPSQQLRSLNGEWR* |
| N= 361 | MKQSTIALALLPLLFTPVTKPRMKQSTIALALLPLLFTPVTKPRTPEMP<br>VLENRAAQGDITAPGGARRLTGDQTAALRDSLSDKPAKNIILLIGDGMG<br>DSEITAARNYAEGAGGFFKGIDALPLTGQYTHYALNKKTGKPDYVTDSA<br>ASATAWSTGVKTYNGALGVDIHEKDHPTILEMAKAAGLATGNVSTAELQ<br>DATPAALVAHVTSRKCYGPSATSEKCPGNALEKGGKGSITEQLLNARAD<br>VTLGGGAKTFAETATAGEWQGKTLREQAQARGYQLVSDAASLNSVTEAN<br>QQKPLLGLFADGNMPVRWLGPKATYHGNIDKPAVTCTPNPQRNDSVPTL<br>AQMTDKAIELSGSGYPYDVPDYAFSTPVWISQAQGIRAGPGSSDKQEGE<br>WPTGLRLSRIGGIHSLAVVLQRRDWENPGVTQLNRLAAHPPFASWRNSE<br>EARTDRPSQQLRSLNGEWR* |
| N= 351 | MKQSTIALALLPLLFTPVTKPRMKQSTIALALLPLLFTPVTKPRTPEMP<br>VLENRAAQGDITAPGGARRLTGDQTAALRDSLSDKPAKNIILLIGDGMG<br>DSEITAARNYAEGAGGFFKGIDALPLTGQYTHYALNKKTGKPDYVTDSA<br>ASATAWSTGVKTYNGALGVDIHEKDHPTILEMAKAAGLATGNVSTAELQ<br>DATPAALVAHVTSRKCYGPSATSEKCPGNALEKGGKGSITEQLLNARAD<br>VTLGGGAKTFAETATAGEWQGKTLREQAQARGYQLVSDAASLNSVTEAN<br>QQKPLLGLFADGNMPVRWLGPKATYHGNIDKPAVTCTPNPQRNDSVPTL |

| | |
|---|---|
| | SGSGYPYDVPDYAFSTPVWISQAQGIRAGPGSSDKQEGEWPTGLRLSRI GGIHSLAVVLQRRDWENPGVTQLNRLAAHPPFASWRNSEEARTDRPSQQ LRSLNGEWR* |
| N= 341 | MKQSTIALALLPLLFTPVTKPRMKQSTIALALLPLLFTPVTKPRTPEMP VLENRAAQGDITAPGGARRLTGDQTAALRDSLSDKPAKNIILLIGDGMG DSEITAARNYAEGAGGFFKGIDALPLTGQYTHYALNKKTGKPDYVTDSA ASATAWSTGVKTYNGALGVDIHEKDHPTILEMAKAAGLATGNVSTAELQ DATPAALVAHVTSRKCYGPSATSEKCPGNALEKGGKGSITEQLLNARAD VTLGGGAKTFAETATAGEWQGKTLREQAQARGYQLVSDAASLNSVTEAN QQKPLLGLFADGNMPVRWLGPKATYHGNIDKPAVTCTPNPSGSGYPYDV PDYAFSTPVWISQAQGIRAGPGSSDKQEGEWPTGLRLSRIGGIHSLAVV LQRRDWENPGVTQLNRLAAHPPFASWRNSEEARTDRPSQQLRSLNGEWR * |
| N= 331 | MKQSTIALALLPLLFTPVTKPRMKQSTIALALLPLLFTPVTKPRTPEMP VLENRAAQGDITAPGGARRLTGDQTAALRDSLSDKPAKNIILLIGDGMG DSEITAARNYAEGAGGFFKGIDALPLTGQYTHYALNKKTGKPDYVTDSA ASATAWSTGVKTYNGALGVDIHEKDHPTILEMAKAAGLATGNVSTAELQ DATPAALVAHVTSRKCYGPSATSEKCPGNALEKGGKGSITEQLLNARAD VTLGGGAKTFAETATAGEWQGKTLREQAQARGYQLVSDAASLNSVTEAN QQKPLLGLFADGNMPVRWLGPKATYHGNISGSGYPYDVPDYAFSTPVWI SQAQGIRAGPGSSDKQEGEWPTGLRLSRIGGIHSLAVVLQRRDWENPGV TQLNRLAAHPPFASWRNSEEARTDRPSQQLRSLNGEWR* |
| N= 321 | MKQSTIALALLPLLFTPVTKPRMKQSTIALALLPLLFTPVTKPRTPEMP VLENRAAQGDITAPGGARRLTGDQTAALRDSLSDKPAKNIILLIGDGMG DSEITAARNYAEGAGGFFKGIDALPLTGQYTHYALNKKTGKPDYVTDSA ASATAWSTGVKTYNGALGVDIHEKDHPTILEMAKAAGLATGNVSTAELQ DATPAALVAHVTSRKCYGPSATSEKCPGNALEKGGKGSITEQLLNARAD VTLGGGAKTFAETATAGEWQGKTLREQAQARGYQLVSDAASLNSVTEAN QQKPLLGLFADGNMPVRWLSGSGYPYDVPDYAFSTPVWISQAQGIRAGP GSSDKQEGEWPTGLRLSRIGGIHSLAVVLQRRDWENPGVTQLNRLAAHP PFASWRNSEEARTDRPSQQLRSLNGEWR* |
| N= 311 | MKQSTIALALLPLLFTPVTKPRMKQSTIALALLPLLFTPVTKPRTPEMP VLENRAAQGDITAPGGARRLTGDQTAALRDSLSDKPAKNIILLIGDGMG DSEITAARNYAEGAGGFFKGIDALPLTGQYTHYALNKKTGKPDYVTDSA ASATAWSTGVKTYNGALGVDIHEKDHPTILEMAKAAGLATGNVSTAELQ DATPAALVAHVTSRKCYGPSATSEKCPGNALEKGGKGSITEQLLNARAD VTLGGGAKTFAETATAGEWQGKTLREQAQARGYQLVSDAASLNSVTEAN QQKPLLGLFSGSGYPYDVPDYAFSTPVWISQAQGIRAGPGSSDKQEGEW PTGLRLSRIGGIHSLAVVLQRRDWENPGVTQLNRLAAHPPFASWRNSEE ARTDRPSQQLRSLNGEWR* |
| N= 307 | MKQSTIALALLPLLFTPVTKPRMKQSTIALALLPLLFTPVTKPRTPEMP VLENRAAQGDITAPGGARRLTGDQTAALRDSLSDKPAKNIILLIGDGMG DSEITAARNYAEGAGGFFKGIDALPLTGQYTHYALNKKTGKPDYVTDSA ASATAWSTGVKTYNGALGVDIHEKDHPTILEMAKAAGLATGNVSTAELQ DATPAALVAHVTSRKCYGPSATSEKCPGNALEKGGKGSITEQLLNARAD VTLGGGAKTFAETATAGEWQGKTLREQAQARGYQLVSDAASLNSVTEAN QQKPLSGSGYPYDVPDYAFSTPVWISQAQGIRAGPGSSDKQEGEWPTGL RLSRIGGIHSLAVVLQRRDWENPGVTQLNRLAAHPPFASWRNSEEARTD RPSQQLRSLNGEWR* |
| N= 305 | MKQSTIALALLPLLFTPVTKPRMKQSTIALALLPLLFTPVTKPRTPEMP VLENRAAQGDITAPGGARRLTGDQTAALRDSLSDKPAKNIILLIGDGMG DSEITAARNYAEGAGGFFKGIDALPLTGQYTHYALNKKTGKPDYVTDSA |

| | |
|---|---|
| | ASATAWSTGVKTYNGALGVDIHEKDHPTILEMAKAAGLATGNVSTAELQ DATPAALVAHVTSRKCYGPSATSEKCPGNALEKGGKGSITEQLLNARAD VTLGGGAKTFAETATAGEWQGKTLREQAQARGYQLVSDAASLNSVTEAN QQKSGSGYPYDVPDYAFSTPVWISQAQGIRAGPGSSDKQEGEWPTGLRL SRIGGIHSLAVVLQRRDWENPGVTQLNRLAAHPPFASWRNSEEARTDRP SQQLRSLNGEWR* |
| N= 301 | MKQSTIALALLPLLFTPVTKPRMKQSTIALALLPLLFTPVTKPRTPEMP VLENRAAQGDITAPGGARRLTGDQTAALRDSLSDKPAKNIILLIGDGMG DSEITAARNYAEGAGGFFKGIDALPLTGQYTHYALNKKTGKPDYVTDSA ASATAWSTGVKTYNGALGVDIHEKDHPTILEMAKAAGLATGNVSTAELQ DATPAALVAHVTSRKCYGPSATSEKCPGNALEKGGKGSITEQLLNARAD VTLGGGAKTFAETATAGEWQGKTLREQAQARGYQLVSDAASLNSVTEAS GSGYPYDVPDYAFSTPVWISQAQGIRAGPGSSDKQEGEWPTGLRLSRIG GIHSLAVVLQRRDWENPGVTQLNRLAAHPPFASWRNSEEARTDRPSQQL RSLNGEWR* |
| N= 291 | MKQSTIALALLPLLFTPVTKPRMKQSTIALALLPLLFTPVTKPRTPEMP VLENRAAQGDITAPGGARRLTGDQTAALRDSLSDKPAKNIILLIGDGMG DSEITAARNYAEGAGGFFKGIDALPLTGQYTHYALNKKTGKPDYVTDSA ASATAWSTGVKTYNGALGVDIHEKDHPTILEMAKAAGLATGNVSTAELQ DATPAALVAHVTSRKCYGPSATSEKCPGNALEKGGKGSITEQLLNARAD VTLGGGAKTFAETATAGEWQGKTLREQAQARGYQLVSDSGSGYPYDVPD YAFSTPVWISQAQGIRAGPGSSDKQEGEWPTGLRLSRIGGIHSLAVVLQ RRDWENPGVTQLNRLAAHPPFASWRNSEEARTDRPSQQLRSLNGEWR* |
| N= 281 | MKQSTIALALLPLLFTPVTKPRMKQSTIALALLPLLFTPVTKPRTPEMP VLENRAAQGDITAPGGARRLTGDQTAALRDSLSDKPAKNIILLIGDGMG DSEITAARNYAEGAGGFFKGIDALPLTGQYTHYALNKKTGKPDYVTDSA ASATAWSTGVKTYNGALGVDIHEKDHPTILEMAKAAGLATGNVSTAELQ DATPAALVAHVTSRKCYGPSATSEKCPGNALEKGGKGSITEQLLNARAD VTLGGGAKTFAETATAGEWQGKTLREQAQASGSGYPYDVPDYAFSTPVW ISQAQGIRAGPGSSDKQEGEWPTGLRLSRIGGIHSLAVVLQRRDWENPG VTQLNRLAAHPPFASWRNSEEARTDRPSQQLRSLNGEWR* |
| N= 271 | MKQSTIALALLPLLFTPVTKPRMKQSTIALALLPLLFTPVTKPRTPEMP VLENRAAQGDITAPGGARRLTGDQTAALRDSLSDKPAKNIILLIGDGMG DSEITAARNYAEGAGGFFKGIDALPLTGQYTHYALNKKTGKPDYVTDSA ASATAWSTGVKTYNGALGVDIHEKDHPTILEMAKAAGLATGNVSTAELQ DATPAALVAHVTSRKCYGPSATSEKCPGNALEKGGKGSITEQLLNARAD VTLGGGAKTFAETATAGESGSGYPYDVPDYAFSTPVWISQAQGIRAGPG SSDKQEGEWPTGLRLSRIGGIHSLAVVLQRRDWENPGVTQLNRLAAHPP FASWRNSEEARTDRPSQQLRSLNGEWR* |
| N= 251 | MKQSTIALALLPLLFTPVTKPRMKQSTIALALLPLLFTPVTKPRTPEMP VLENRAAQGDITAPGGARRLTGDQTAALRDSLSDKPAKNIILLIGDGMG DSEITAARNYAEGAGGFFKGIDALPLTGQYTHYALNKKTGKPDYVTDSA ASATAWSTGVKTYNGALGVDIHEKDHPTILEMAKAAGLATGNVSTAELQ DATPAALVAHVTSRKCYGPSATSEKCPGNALEKGGKGSITEQLLNARSG SGYPYDVPDYAFSTPVWISQAQGIRAGPGSSDKQEGEWPTGLRLSRIGG IHSLAVVLQRRDWENPGVTQLNRLAAHPPFASWRNSEEARTDRPSQQLR SLNGEWR* |
| N= 241 | MKQSTIALALLPLLFTPVTKPRMKQSTIALALLPLLFTPVTKPRTPEMP VLENRAAQGDITAPGGARRLTGDQTAALRDSLSDKPAKNIILLIGDGMG DSEITAARNYAEGAGGFFKGIDALPLTGQYTHYALNKKTGKPDYVTDSA ASATAWSTGVKTYNGALGVDIHEKDHPTILEMAKAAGLATGNVSTAELQ DATPAALVAHVTSRKCYGPSATSEKCPGNALEKGGKGSGSGYPYDVPDY |

| | |
|---|---|
| | AFSTPVWISQAQGIRAGPGSSDKQEGEWPTGLRLSRIGGIHSLAVVLQR<br>RDWENPGVTQLNRLAAHPPFASWRNSEEARTDRPSQQLRSLNGEWR* |
| N= 141 | MKQSTIALALLPLLFTPVTKPRMKQSTIALALLPLLFTPVTKPRTPEMP<br>VLENRAAQGDITAPGGARRLTGDQTAALRDSLSDKPAKNIILLIGDGMG<br>DSEITAARNYAEGAGGFFKGIDALPLTGQYTHYALSGSGYPYDVPDYAF<br>STPVWISQAQGIRAGPGSSDKQEGEWPTGLRLSRIGGIHSLAVVLQRRD<br>WENPGVTQLNRLAAHPPFASWRNSEEARTDRPSQQLRSLNGEWR* |

### List of Primers

| Construct Name | Forward Primer 5' → 3' | Reverse Primer 5' → 3' |
|---|---|---|
| *N*= 491 | TCAGGATCGGGCTACCC | TAGCCCGATCCTGAGAAGAGATCGGTCTGGTCG |
| *N*= 471 | TCAGGATCGGGCTACCC | TAGCCCGATCCTGAAATACGCAACTGACTGCCG |
| *N*= 461 | TCAGGATCGGGCTACCC | TAGCCCGATCCTGATGAATCCTCTTCGGAGTTCCC |
| *N*= 450 | TCAGGATCGGGCTACCCATACG | CATCACTGCGCCATCTTTGGTATTTAGC |
| *N*= 440 | TCAGGATCGGGCTACCCATACG | CTGGGTGAGGCCCGGAG |
| *N*= 435 | TCAGGATCGGGCTACCCATACG | AGCTTTGGTATCCGGCGCAAC |
| *N*= 431 | TCAGGATCGGGCTACCC | TAGCCCGATCCTGACGGCGCAACAATCTGG |
| *N*= 426 | TCAGGATCGGGCTACCCATACG | GCTGGCGTGGGCGTGATC |
| *N*= 421 | TCAGGATCGGGCTACCC | TAGCCCGATCCTGAATCAGCGGTGACTATGACCAG |
| *N*= 415 | TCAGGATCGGGCTACCCATACG | CAGCGTGTTACCCTCCTTTTTAGC |
| *N*= 411 | TCAGGATCGGGCTACCC | TAGCCCGATCCTGACTCCTTTTTAGCGAATTCCAGCG |
| *N*= 409 | TCAGGATCGGGCTACCCATACG | TTTAGCGAATTCCAGCGCCCG |
| *N*= 407 | TCAGGATCGGGCTACCCATACG | GAATTCCAGCGCCCGTTGTACG |
| *N*= 403 | TCAGGATCGGGCTACCCATACG | CCGTTGTACGGCTTCATCGAGATC |
| *N*= 401 | TCAGGATCGGGCTACCC | TAGCCCGATCCTGATACGGCTTCATCGAGATCGAC |
| *N*= 391 | TCAGGATCGGGCTACCC | TAGCCCGATCCTGACTGTTTATCGATTGACGCACCTTC |
| *N*= 381 | TCAGGATCGGGCTACCC | TAGCCCGATCCTGACAGGAAAAAGCCTTTCTCATTTTTACTCAAC |
| *N*= 371 | TCAGGATCGGGCTACCC | TAGCCCGATCCTGACAATTCAATGGCTTTGTCGGTC |
| *N*= 361 | TCAGGATCGGGCTACCC | TAGCCCGATCCTGACAGGGTTGGTACACTGTCATTAC |
| *N*= 351 | TCAGGATCGGGCTACCC | TAGCCCGATCCTGAATTTGGCGTACAGGTGACTG |
| *N*= 341 | TCAGGATCGGGCTACCC | TAGCCCGATCCTGAGATATTGCCATGGTACGTTGCTTTC |
| *N*= 331 | TCAGGATCGGGCTACCC | TAGCCCGATCCTGATAGCCAGCGCACTGG |
| *N*= 321 | TCAGGATCGGGCTACCC | TAGCCCGATCCTGAAAACAGGCCAAGCAGGGG |
| *N*= 311 | TCAGGATCGGGCTACCC | TAGCCCGATCCTGATAGCCAGCGCACTGG |
| *N*= 307 | TCAGGATCGGGCTACCCATACG | CAGGGGTTTTTGCTGATTCGCTTC |
| *N*= 301 | TCAGGATCGGGCTACCC | TAGCCCGATCCTGAATCGCTCACCAACTGATAACC |
| *N*= 291 | TCAGGATCGGGCTACCC | TAGCCCGATCCTGATGCCTGTTCACGC |

| | | |
|---|---|---|
| *N*= 281 | TCAGGATCGGGCTACCC | TAGCCCGATCCTGATTCACCAGCGGTTGCC |
| *N*= 271 | TCAGGATCGGGCTACCC | TAGCCCGATCCTGATTTTGCGCCG |
| *N*= 251 | TCAGGATCGGGCTACCC | TAGCCCGATCCTGATCCTTTTCCGCCTTTTTCCAG |
| *N*= 241 | TCAGGATCGGGCTACCC | TAGCCCGATCCTGACGGACATTTTTCACTGGTCGCGCTCG |
| *N*= 231 | TCAGGATCGGGCTACCC | TAGCCCGATCCTGAGTAGCATTTGCGCGAGG |
| *N*= 141 | TCAGGATCGGGCTACCC | TAGCCCGATCCTGACAGCGCATAGTGAGTGTATTGC |
| C190A | GCATACGGTCCGAGCGCGA | TTTGCGCGAGGTCACATGTGC |
| C200A | GCACCGGGTAACGCTCTGGAAA | TTTTTCACTGGTCGCGCTCGGA |
| C308A | GCAACGCCAAATCCGCAACGTAAT | GGTGACTGCGGGCTTATCGATAT |
| C358A | GCAGGGCAAATTGGCGAGACG | AGGATTCGCAGCATGATCCTGTTTATC |
